# ProtEnrich: Residual Multimodal Enrichment of Protein Sequence Embeddings

**DOI:** 10.64898/2026.08.24.746797

**Authors:** Gabriel Bianchin de Oliveira, Fahad Saeed

## Abstract

Protein language models effectively capture evolutionary and functional signals from sequence data but lack explicit representation of the biophysical properties that govern protein structure and dynamics. Existing multimodal approaches attempt to integrate such physical information through direct fusion, often requiring multimodal inputs at inference time and distorting the geometry of the sequence embedding space, which can disrupt the semantic organization learned from evolutionary information. Consequently, a fundamental challenge of how to incorporate structural and dynamical knowledge into sequence representations without disrupting their semantic organization, enabling sequence-based models to better capture the biophysical properties governing protein structure and function. We introduce ProtEnrich, a representation learning framework based on a residual multimodal enrichment paradigm. ProtEnrich decomposes sequence embeddings into two complementary latent subspaces, an anchor subspace that preserves sequence semantics, and an alignment subspace that encodes biophysical relationships. By converting multimodal information derived from ProstT5 and RocketSHP to a low-energy residual component, our approach injects physical representation while maintaining the original sequence embedding while preserving their original semantic geometry, avoiding the need for multimodal inputs at inference time. Across eight diverse protein foundational models trained on 550,120 SwissProt proteins with AlphaFold structures, enriched embeddings improved zero-shot remote homology retrieval, increasing Precision@10 and MRR by up to 0.13 and 0.11, respectively. Downstream performance also improved on structure-dependent tasks, reducing fluorescence prediction error by up to 16% and increasing metal ion binding AUCROC by up to 2.4 points, while requiring only sequence input at inference. Source code is available at https://github.com/pcdslab/ProtEnrich, pretrained models and datasets are available at https://huggingface.co/collections/SaeedLab/protenrich.

## 1 Introduction

Protein language models have transformed computational biology by enabling large-scale learning of protein representations directly from sequence data. These models capture evolutionary signals through self-supervised training, achieving remarkable performance across diverse downstream tasks [1–3]. While useful, in real-world, protein properties are determined by three-dimensional fold and conformational flexibility, which emerge from physical interactions between amino acids. While recent advances in structure prediction have significantly expanded the availability of structural data [4–6], most proteins still lack experimentally validated structures, and predicted models typically represent static conformations that do not capture dynamic behavior. Consequently, large-scale protein analysis remains predominantly sequence-driven, creating a fundamental gap between the physical reality of proteins and their computational representations. Despite their success, protein language models representations also remain limited to sequence and lack explicit encoding of the biophysical principles governing protein structure and dynamics, potentially limiting their ability to capture structure and dynamics-dependent protein properties.

Existing multimodal approaches attempt to bridge this gap by integrating sequence, structure, and other modalities into unified representations [7–10]. However, these methods typically require multimodal inputs at inference time [7, 10] or rely on fusion strategies that alter the geometry of the original sequence embedding space [8, 9]. Such processes may introduce representation distortion, potentially disrupting the semantic organization learned from evolutionary data and affecting performance in tasks primarily driven by sequence information. Thus, a central challenge of incorporating biophysical properties into sequence representations without compromising their intrinsic semantic structure, remains unsolved.

In this work, we introduce ProtEnrich, a framework based on a residual multimodal enrichment paradigm that transfers structural and dynamical knowledge into sequence embeddings while preserving their original representation. ProtEnrich decomposes sequence representations into two complementary latent subspaces, an anchor subspace that maintains sequence semantics, and a multimodal alignment subspace that learns to encode biophysical relationships through geometric alignment with structural and dynamical embeddings. By limiting multimodal information to a low-energy residual subspace, our approach enables the injection of physical properties without distorting the original embedding. To create this alignment between sequences and physical properties, ProtEnrich uses pretrained structural representations derived from ProstT5 [11] and dynamical descriptors from RocketSHP [12] to capture conformational flexibility.

Our main contributions are:

- We introduce a residual multimodal enrichment paradigm that enables the transfer of structural and dynamical properties into protein sequence representations without requiring multimodal inputs at inference time, enabling sequence-based models to capture biophysical information relevant to protein properties prediction.
- We demonstrate that protein sequence embeddings can be decomposed into orthogonal latent subspaces separating semantic sequence information from biophysical signals, enabling controlled integration of structural and dynamical information while preserving the semantic organization of the original embedding space.
- We show that multimodal knowledge can be injected as a low-energy residual component, preserving the geometry of the original sequence embedding manifold and enabling multimodal enrichment without degrading the semantic space learned from sequence data.

We evaluate ProtEnrich across eight protein sequence foundational models ranging from diverse architecture paradigms. The enriched representations consistently improve zero-shot structural retrieval, latent feature imputation, and downstream predictive tasks while requiring only sequence information at inference time. ProtEnrich’s embeddings improved zero-shot remote homology retrieval, increasing Precision@10 and MRR by up to 0.13 and 0.11, respectively, and reduced the error by up to 16% in regression tasks compared to the original protein sequence models embeddings. Furthermore, our analyses reveal that the learned multimodal alignment subspace preserves structural, and dynamical similarity relationships and forms orthogonal latent representation with anchor sequence-based signals, providing new insights into the organization of protein representation spaces.

## 2 Methods

### 2.1 Problem Formulation

Protein language models produce embeddings that capture sequence information but lack explicit representations of structural and dynamical properties that influence protein behavior. Our goal is to incorporate biophysical information into sequence representations while preserving their semantic organization and requiring only sequence input at inference time.

The problem is formulated as the residual multimodal enrichment of protein sequence representations with structural and dynamical properties, relying exclusively on sequence inputs during inference. This requires transferring multimodal knowledge into a sequence latent space without disrupting the semantic information encoded by the original sequence embedding. Formally, given a sequence embedding *x_seq_*obtained from a protein foundational model and auxiliary structural *x_str_* and dynamical *x_dyn_* embeddings, our goal is to learn an enriched representation *h_enrich_* that incorporates residual multimodal information during training but depends solely on sequence input during inference.

### 2.2 ProtEnrich

Figure 1 illustrates the ProtEnrich architecture used during the pretraining stage. ProtEnrich adopts a dual latent subspace framework that learns two complementary representations from the sequence embedding. The first representation, referred to as the anchor representation *h_anchor_*, preserves the original sequence semantics and is optimized to retain the information contained in the input sequence embedding. The second representation, referred to as the multimodal alignment representation *h_algn_*, is designed to capture structural and dynamical relationships. This representation is trained to align geometrically with structural and dynamical embeddings, enabling the transfer of multimodal knowledge into the sequence domain.

**Figure 1:**
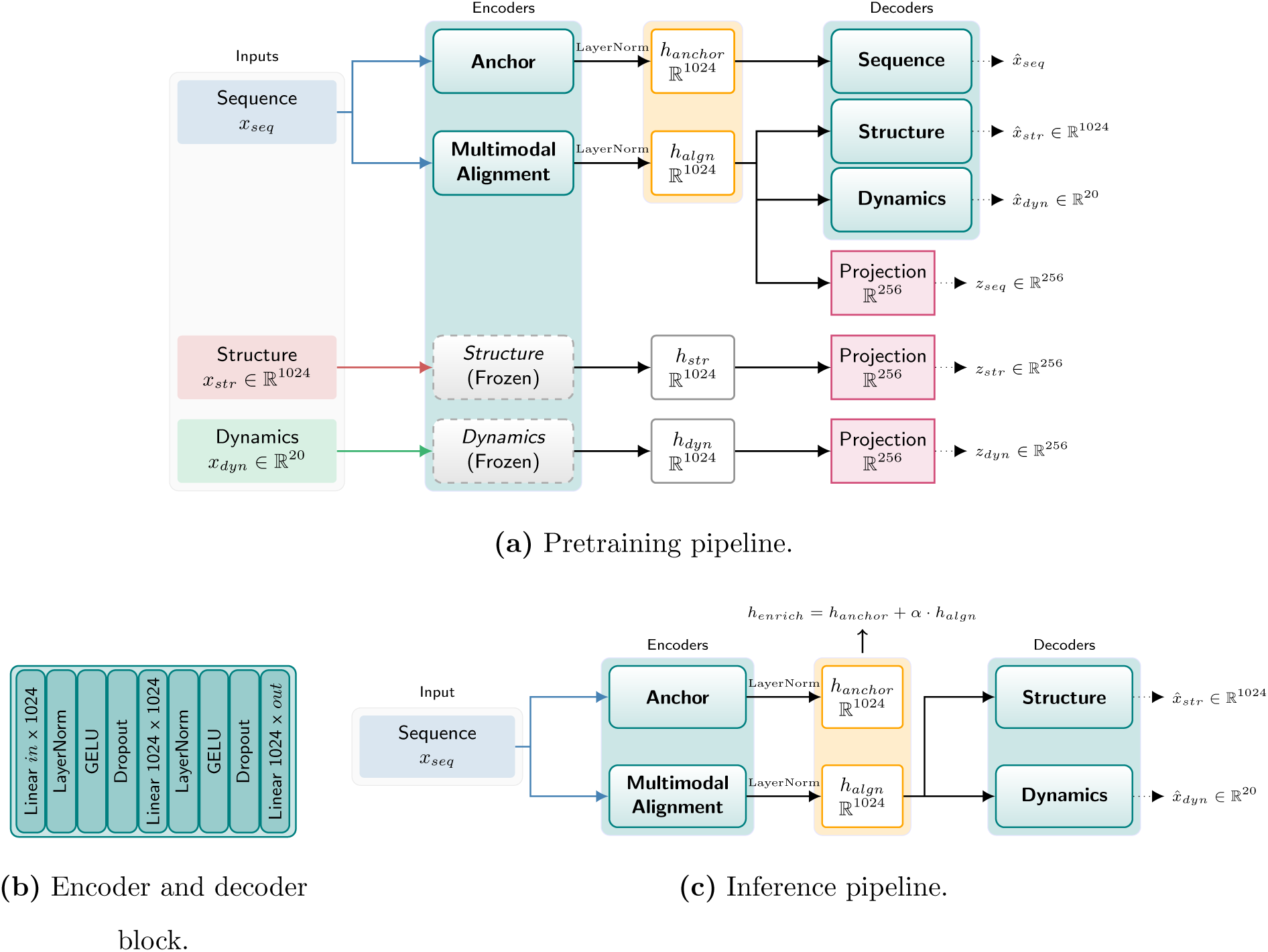
Overview of the ProtEnrich framework. (a) Pretraining pipeline. During pretraining, a frozen protein sequence embedding obtained from a foundational protein language model is processed by two parallel encoders that learn complementary latent representations. In parallel, pretrained structural embeddings and dynamical embeddings are projected into the same latent space through frozen encoders. All representations are mapped into a shared 1024-dimensional latent space. The multimodal alignment representation is optimized to align geometrically with both structural and dynamical modalities, while reconstruction decoders enforce recovery of original embeddings. (b) Encoder and decoder block. Each encoder and decoder consists of a multi-layer perceptron that projects input embeddings into a specific output dimension (c) Inference pipeline. At inference time, only the sequence embedding is required. The anchor and alignment encoders generate *h_anchor_* and *h_algn_*, which are combined through a controllable residual mechanism to generate *h_enrich_*. *h_algn_*is also capable to generate latent embedding for structural and dynamical information.

Both representations are learned jointly through separate encoders applied to the sequence embedding. Structural *h_str_* and dynamical *h_dyn_* embeddings are projected into the same latent space through frozen encoders, ensuring that they serve as reference modalities rather than trainable components. All representations are mapped to a shared continuous latent space, such that {*h_anchor_, h_algn_, h_str_, h_dyn_*} ∈ ℝ^1024^. This latent space dimension was chosen to balance expressive capacity and computational efficiency.

This dual-subspace design enables specialization of latent signals, allowing sequence semantics and aligned multimodal enrichment to be encoded separately while maintaining compatibility between them. Figure 1 presents the encoder architecture used for the anchor, alignment, structural, and dynamical embeddings.

### 2.3 Pretraining Objectives

The pretraining of ProtEnrich is performed by jointly minimizing three complementary objectives: (i) geometric alignment between modalities, (ii) reconstruction of original embeddings, and (iii) energy regularization of the enrichment representation.

#### 2.3.1 Alignment Objective

To transfer structural and dynamical information into the multimodal alignment representation, we employ a contrastive alignment objective based on the InfoNCE loss [13]. All modalities are projected into a lower-dimensional contrastive space of dimension 256, where *z_algn_* denotes the projected alignment representation, *z_str_* denotes the projected structural embedding, and *z_dyn_*denotes the projected dynamical embedding.

The alignment objective maximizes the cosine similarity between matched sequence-structure and sequence-dynamics pairs within a batch, while minimizing the similarity with mismatched examples. To dynamically scale the logits and optimize the contrastive landscape during training, we introduce a learnable temperature parameter *τ*, initialized at 0.1. Equation 1 presents the InfoNCE loss, where *N* is the batch size, sim(*u, v*) denotes the cosine similarity, and *z_aux_* is a modality, where *aux* ∈ {*str, dyn*}.

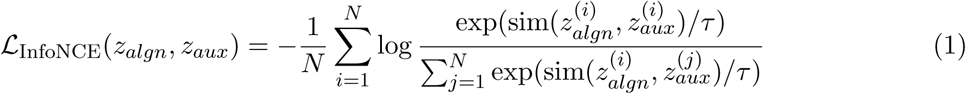

The overall contrastive alignment loss is the sum of the objectives for both structural and dynamical modalities, as defined in Equation 2.

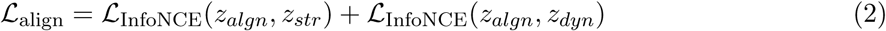

#### 2.3.2 Reconstruction Objectives

To ensure that the alignment representation encodes meaningful multimodal information rather than trivial correlations, we introduce reconstruction objectives using Mean Squared Error (MSE). Structural and dynamical embeddings are reconstructed from the multimodal alignment representation *h_algn_* using dedicated decoders, defined in Figure 1, enforcing that this latent space retains sufficient information to recover the original modalities. Let *x̂_str_* and *x̂_dyn_* denote the reconstructed structural and dynamical embeddings, respectively. Given a batch of size *N*, the multimodal reconstruction losses are defined as equations 3 and 4 for reconstruction of structural and dynamical information, respectively.

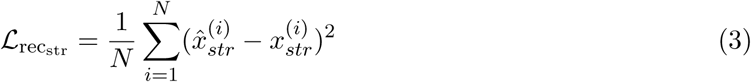

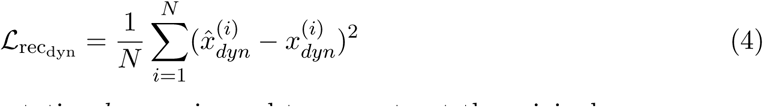

In addition, the anchor representation *h_anchor_* is used to reconstruct the original sequence embedding *x_seq_*via a sequence decoder, yielding the reconstruction *x̂_seq_*. This objective prevents the model from losing sequence semantics during pretraining and ensures that the anchor subspace preserves the original information content. The sequence reconstruction loss is formally given by Equation 5.

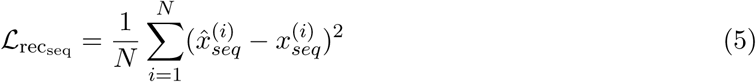

#### 2.3.3 Energy Regularization

Since structural and dynamical information should act as complementary signals rather than dominate the sequence representation, we apply an energy regularization term on the alignment representation. This term minimizes the squared *ℓ*_2_ norm of the alignment embedding, encouraging multimodal enrichment to behave as a low-energy residual signal injected into the sequence representation. Equation 6 presents the energy-based loss.

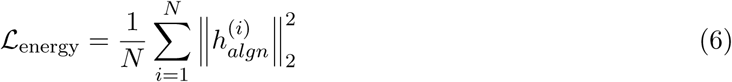

#### 2.3.4 Pretraining Loss

The final training loss combines contrastive alignment, reconstruction losses, and energy regularization, defined in Equation 7.

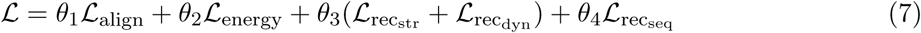

The weighting coefficients were selected based on preliminary sensitivity analyses to balance alignment strength and residual magnitude control. Specifically, equal weights were assigned to alignment and energy terms to balance multimodal knowledge transfer and residual control, with *θ*_1_ and *θ*_2_ equal to 1.0. Reconstruction losses for structural and dynamical embeddings were assigned lower weights to act as auxiliary constraints, with *θ*_3_ equal to 0.5, while sequence reconstruction was weighted minimally to ensure preservation of sequence information, with *θ*_4_ equal to 0.1.

### 2.4 Controllable Residual Enrichment Mechanism

The final enriched representation *h_enrich_* is obtained by combining the anchor *h_anchor_* and multimodal alignment *h_algn_* representations through a controllable mixing mechanism. A coefficient *α* regulates the contribution of the alignment representation to the final embedding, as presented in Equation 8. The coefficient *α* is initialized to 0.039 based on preliminary validation experiments and kept fixed during the experiments reported in this study. When fine-tuning is performed, *α* is allowed to be learned jointly with downstream parameters, enabling task-specific adjustment of the residual strength.

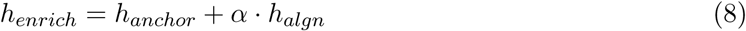

### 2.5 Experimental Setup

#### 2.5.1 Pretraining Data

To pretrain the ProtEnrich framework, we curated a dataset comprising 550,120 protein samples from the SwissProt database [14]. To ensure high-quality structural alignment, the dataset was rigorously filtered to include only proteins with available three-dimensional structures in the AlphaFoldDB [15]. The pretraining phase was executed across all eight sequence foundational model variations for 30 epochs using 95% of the data for training and 5% for validation.

#### 2.5.2 Embedding Representation Models

For structural representations, we utilized the encoder from ProstT5 [11]. This model encodes the 3D structure using the FoldSeek [16] alphabet, outputting a 1024-dimensional continuous vector for each protein by averaging the token representations. Considering the dynamical representations, we employed RocketSHP [12]. By leveraging sequence and structural files in FASTA and PDB, respectively, RocketSHP captures global structural changes via the FoldSeek dictionary, yielding a compact 20-dimensional embedding.

To evaluate the model-agnostic nature of ProtEnrich, we benchmarked eight diverse protein sequence models encompassing encoder, decoder, encoder-decoder, and convolutional architectures. Specifically, our evaluation includes four encoder-based models (ProtBERT [17], ESM Cambrian 600M [18], ESM2 T36 [6], and ESM1b [19]), two encoder-decoder models (ProtT5 [17] and Ankh 3 XL [20], from which we utilized only their encoder components), one decoder-based model (ProGen2 [21]), and one convolutional architecture (CARP 640M [22]). For all sequence models, the final protein representation was obtained by computing the mean of the amino acid tokens, with sequences truncated to a maximum length of 1022 amino acids. Regarding the representation size of each model, ProtBERT and ProtT5 encode the protein sequence using 1024-dimensional vectors, while ESM1b and CARP 640M utilize 1280 dimensions, ProGen2 uses 1536, ESM Cambrian 600M operates with 1156, and Ankh 3 XL employs a 2560-dimensional space. All pretrained sequence models were used in frozen mode without fine-tuning.

#### 2.5.3 Evaluation Protocol

We systematically evaluated ProtEnrich against baseline sequence protein language models across three distinct settings, i.e., zero-shot retrieval, latent feature imputation, and downstream predictive tasks.

For zero-shot retrieval, we utilized two benchmarks. The first is the TAPE [23] remote homology dataset, where the goal is to retrieve proteins sharing the same fold, superfamily, or family. The second is an out-of-distribution set of 10,000 TrEMBL proteins, strictly held out during pretraining, that possess available AlphaFoldDB structures. Within this hold-out set, we defined relevant targets as the top 1% of proteins exhibiting the highest structural and dynamical embedding similarities. Retrieval performance was quantified using Precision at *K*, Recall at *K*, Accuracy at *K*, and Mean Reciprocal Rank (MRR), defined in equations 9, 10, 11, and 12. In the equations, *N* denote the number of query proteins, *i* indicate a query, *Rel_i,k_* is a binary indicator equal to 1 if the protein at rank *k* is relevant and 0 otherwise, *R_i_* denote the total number of relevant proteins for query *i*, **1**(·) denote the indicator function, and *rank_i_*denote the rank position of the first relevant protein. In our experiments, we adopted *K* = 10.

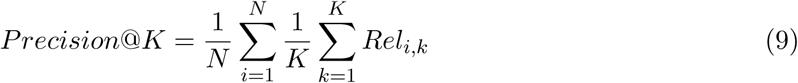

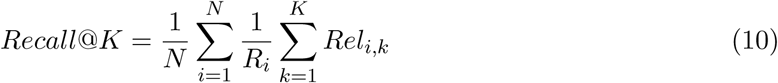

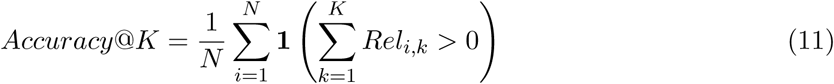

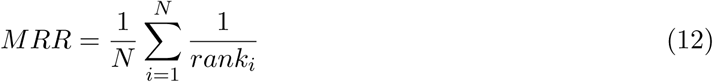

Next, we assessed the model’s ability to impute structural and dynamical features exclusively from sequence inputs. On the 10,000 TrEMBL hold-out proteins, we computed the mean cosine similarity between the imputed embeddings and the ground-truth modalities, comparing these scores against a random permutation baseline. Furthermore, we evaluated the practical utility of these imputed features on the metal ion binding dataset, a standard benchmark from the literature. To ensure a reliable ground truth, this dataset was strictly filtered to include only proteins with available AlphaFold structures. We benchmarked the sequence-derived imputed representations against the true multimodal embeddings to determine whether the injected physical information yielded predictive gains. These evaluations were performed using a 2-layer multi-layer perceptron with 1024 neurons. Hyperparameters were optimized via Optuna [24] over 100 trials, with Area Under Curve ROC (AUCROC) results averaged across 10 random seeds.

Finally, we comprehensively benchmarked the enriched embeddings *h_enrich_* against the original sequence embeddings *h_seq_* for all each protein sequence foundational models on various downstream predictive tasks. These spanned regression (thermostability [25] and fluorescence [23]) and diverse classification benchmarks, including binary (subcellular localization [26], metal ion binding [27], protein-protein interaction [28], and solubility [29]), multiclass (localization [26]), and multilabel (protein function prediction) tasks. Full dataset details are provided in the Supplementary Material. Consistent with the imputation experiments, we trained a 2-layer multi-layer perceptron optimized with Optuna and averaged over 10 independent runs. For the PPI task specifically, inputs were constructed by concatenating the sequence vectors of the interacting protein pair. For downstream evaluation, regression tasks were assessed using Root Mean Squared Error (RMSE), binary classification tasks using AUCROC, multiclass classification using Balanced Accuracy, and multilabel protein function prediction using weighted *F*_max_ (*wF*_max_) [30]. All downstream models were trained using the AdamW [31] optimizer with early stopping based on validation performance. Learning rates were searched in the range from 0.00001 up to 0.005, dropout in range from 0.0 to 0.5, and batch size in {32, 64, 128}.

## 3 Results and Discussion

### 3.1 Zero-Shot Retrieval

We first evaluated ProtEnrich on zero-shot retrieval using the remote homology benchmark. Figure 2 reports the performance delta between enrich and original embeddings, highlighting the effect of residual multimodal enrichment over the original sequence embeddings across fold, superfamily, and family levels. Complete absolute results for all metrics and models are provided in the Supplementary Material. Across all evaluated models and hierarchical levels, ProtEnrich consistently improved retrieval performance relative to the original sequence embeddings. Improvements were observed at the fold, superfamily, and family levels, indicating that the enrichment mechanism enhances both local and remote structural sensitivity. Notably, the largest improvements were observed for ProtBERT. At the family level, Precision@10 increased from 0.369 to 0.490, Recall@10 from 0.204 to 0.294, Accuracy@10 from 0.808 to 0.901, and MRR from 0.733 to 0.846.

**Figure 2:**
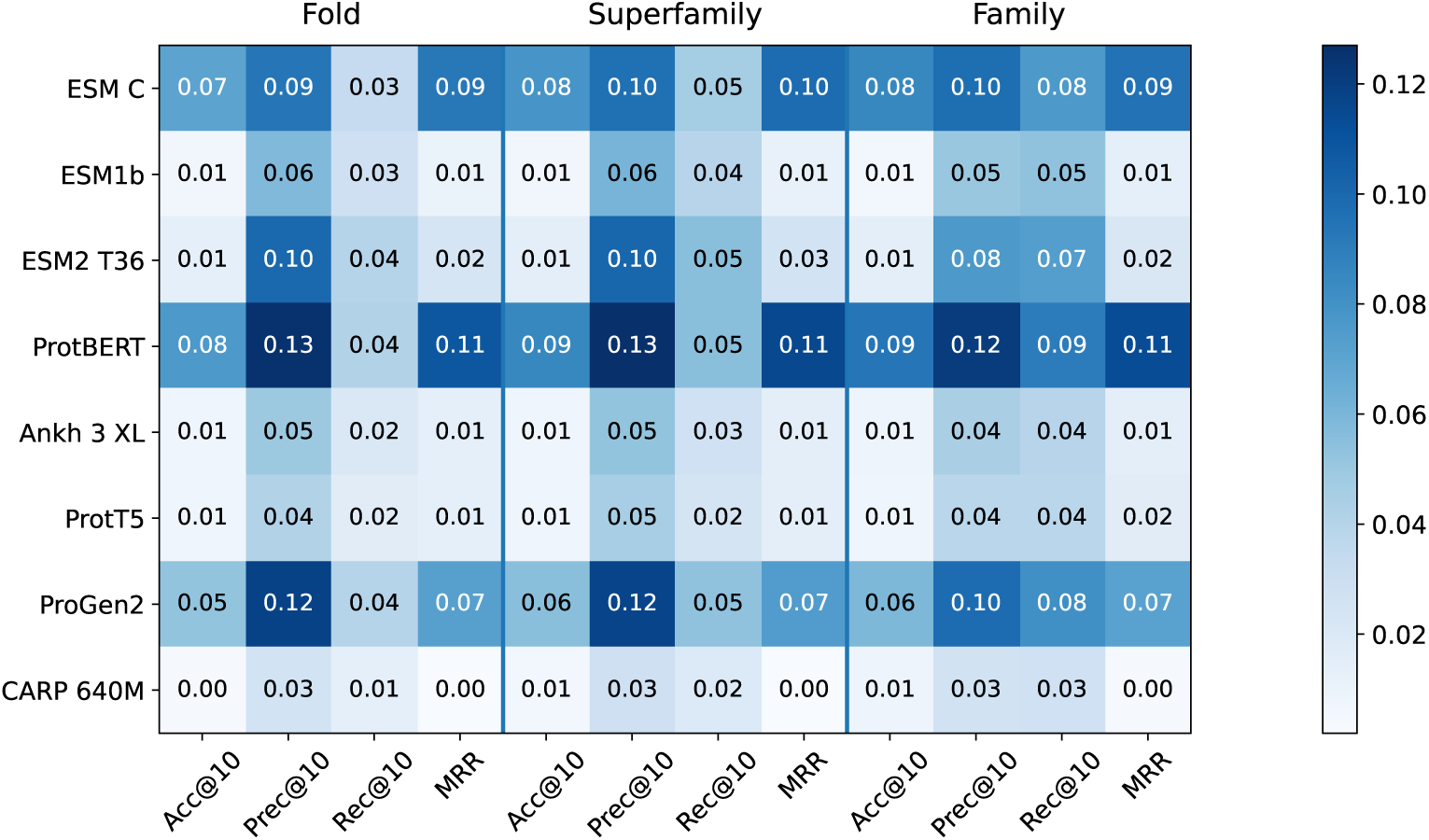
Performance delta between ProtEnrich and original embeddings on the remote homology benchmark across fold, superfamily, and family levels.

Next, we evaluated zero-shot retrieval on an out-of-distribution set of 10,000 TrEMBL proteins, defining relevance based on structural and dynamical similarity. The results are presented in the Supplementary Material. In most cases, enriched embeddings outperformed the original sequence embeddings. For encoder-decoder-based architectures (Ankh 3 XL and ProtT5), the enrichment yielded comparable but slightly lower performance, suggesting that this kind of model may already encode implicit structural and dynamical information from sequence alone.

Together, these results indicate that residual multimodal enrichment reshapes the embedding geometry to enhance structural and dynamical sensitivity, even without task-specific training or multimodal input at inference time. These findings are particularly relevant for remote homology detection, where proteins may share conserved folds despite low sequence identity. The highest gains were obtained using ProtBERT embeddings. While other models also exhibited consistent gains, the magnitude of improvement varied across architectures. This suggests that ProtEnrich acts as a geometry regularizer whose impact depends on the baseline structural organization of the underlying embedding space. Models with weaker initial structural alignment benefit more substantially from residual multimodal enrichment.

### 3.2 Latent Feature Imputation

Motivated by the improved structural sensitivity observed in zero-shot retrieval, we next evaluated whether the alignment subspace can reconstruct structural and dynamical representations directly from sequence. First, we computed the mean cosine similarity between generated embeddings and their corresponding ground-truth modalities on the 10,000 out-of-distribution proteins. As a baseline, we measured similarity between generated embeddings and randomly permuted structural or dynamical embeddings. The results are presented in Table 1. Generated embeddings exhibited substantially higher cosine similarity to their ground-truth counterparts than to the permuted baseline across all models, suggesting that the alignment subspace captures modality-specific information beyond trivial regression.

**Table 1:** Mean cosine similarity (± standard deviation) between generated embeddings and their ground-truth structural or dynamical representations (Orig-Gen), and between generated embeddings and randomly permuted modalities (Rand-Gen), computed over 10,000 out-of-distribution proteins. Higher values indicate greater similarity.

| Model | Structure Similarity |  | Dynamics Similarity |  |
| --- | --- | --- | --- | --- |
|  | Orig-Gen | Rand-Gen | Orig-Gen | Rand-Gen |
| ESM C 600M | 0.946 ± 0.037 | 0.652 ± 0.151 | 0.994 ± 0.013 | 0.817 ± 0.196 |
| ESM1b | 0.949 ± 0.037 | 0.650 ± 0.149 | 0.993 ± 0.013 | 0.806 ± 0.206 |
| ESM2 T36 | 0.948 ± 0.036 | 0.659 ± 0.146 | 0.995 ± 0.011 | 0.818 ± 0.195 |
| ProtBERT | 0.937 ± 0.040 | 0.664 ± 0.142 | 0.993 ± 0.013 | 0.818 ± 0.198 |
| Ankh | 0.944 ± 0.033 | 0.666 ± 0.140 | 0.994 ± 0.011 | 0.827 ± 0.188 |
| ProtT5 | 0.957 ± 0.032 | 0.650 ± 0.148 | 0.994 ± 0.012 | 0.807 ± 0.204 |
| ProGen2 | 0.923 ± 0.045 | 0.677 ± 0.139 | 0.990 ± 0.016 | 0.817 ± 0.197 |
| CARP 640M | 0.945 ± 0.036 | 0.659 ± 0.148 | 0.994 ± 0.013 | 0.816 ± 0.201 |

Then, we assessed the practical utility of these generated features on the metal ion binding task. We evaluated multiple combinations of original and generated structural and dynamical embeddings (see Supplementary Material for full details). Across models, generated features consistently improved performance over the sequence-only baseline in at least one configuration. In most cases, performance followed a consistent hierarchy, where sequence-only embeddings yielded the lowest performance, sequence combined with generated features achieved intermediate performance, and sequence combined with true structural and dynamical embeddings achieved the highest performance. This pattern aligns with the residual design principle, indicating that generated embeddings approximate, but do not fully replace, the information content of true structural modalities. The largest improvements were observed for ESM Cambrian 600M, which showed an increase of 3.8 of AUCROC when concatenating generated structural and dynamical features.

### 3.3 Downstream Predictive Tasks

We next evaluated enriched representations *h_enrich_* on a diverse set of downstream regression and classification tasks. For each protein language model, we compared the original frozen sequence embedding against *h_enrich_*, which was computed using sequence input only.

Results are reported in Table 2. Enriched embeddings yielded notable improvements in fluorescence prediction and metal ion binding. For ProGen2, fluorescence error decreased by approximately 16%, while for ESM2 T36, metal ion binding performance increased by 2.4 AUCROC points. These tasks are known to depend strongly on structural and interaction-related properties, and may therefore benefit from increased structural sensitivity introduced by residual enrichment. For other tasks, such as solubility, performance differences were minimal or slightly negative. This suggests that sequence-dominated tasks may not benefit substantially from structural enrichment, reinforcing the importance of controlled residual scaling to preserve sequence semantics.

**Table 2:**
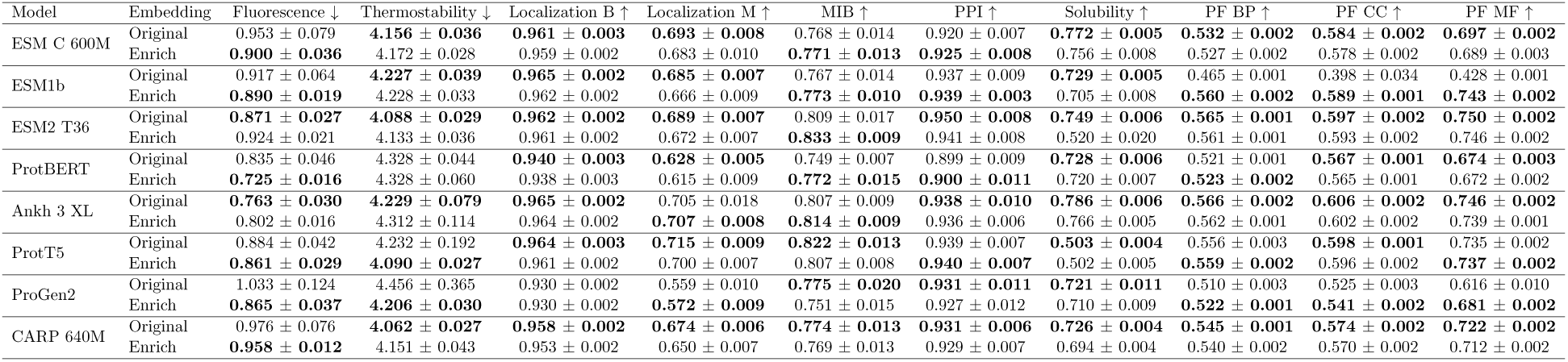
Downstream performance (mean ± standard deviation) for the original and enriched embeddings. The best result within each model is shown in bold. Localization B and Localization M denote binary and multiclass localization, respectively. MIB refers to metal ion binding, and PPI to protein-protein interaction. PF stands for protein function prediction and is divided into three Gene Ontology categories: BP (Biological Process), CC (Cellular Component), and MF (Molecular Function). For regression tasks, lower values indicate better performance, whereas for classification tasks, higher values indicate better performance.

### 3.4 Comparison with Structure-Aware Pretraining

To contextualize ProtEnrich with respect to recent structure-aware protein language models, we compared it against ISM [9], a structure-informed variant of ESM2 that integrates structural supervision during backbone pretraining while requiring only sequence input at inference time. As ISM is specifically designed for ESM2 architectures, we performed the comparison using the ESM2 T36 backbone.

We evaluated both approaches on structure-relevant benchmarks, including zero-shot remote homology retrieval (family, superfamily, and fold levels), zero-shot structural similarity retrieval, and downstream tasks with strong structural dependence (metal ion binding and fluorescence prediction). Table 3 reports the results for family-level remote homology and metal ion binding, while complete results are provided in the Supplementary Material. ProtEnrich consistently improves over the original ESM2 T36 embeddings and outperforms ISM on family-level remote homology retrieval, particularly in Precision@10 and Recall@10. In addition, ProtEnrich achieves the highest performance on metal ion binding prediction, indicating improved structure-aware functional discrimination without modifying the backbone architecture. In contrast, ISM shows gains in pure structural similarity retrieval (in Supplementary Material), reflecting the direct incorporation of structural supervision during large-scale backbone pretraining.

**Table 3:** Comparison between ProtEnrich and ISM on structure-sensitive tasks using the ESM2 T36 backbone. Remote homology results are reported at the family level. Metal ion binding (MIB) is evaluated using mean ± standard deviation across runs. Higher values indicate better performance.

| Model | Remote Homology - Family | | | | MIB ↑ |
| --- | --- | --- | --- | --- | --- |
|  | A@10 | P@10 | R@10 | MRR |  |
| Original | 0.963 | 0.634 | 0.429 | 0.937 | $0.809 \pm 0.017$ |
| ProtEnrich (Ours) | **0.977** | **0.710** | **0.502** | **0.958** | **$0.833 \pm 0.009$** |
| ISM | 0.966 | 0.639 | 0.436 | 0.944 | $0.821 \pm 0.008$ |

Overall, these results suggest that backbone-level structural pretraining primarily enhances explicit structural similarity, whereas residual multimodal enrichment more effectively improves fold-level generalization and structure-informed functional prediction without requiring architectural modification.

### 3.5 Ablation Study

To better characterize the functional roles of the learned latent subspaces, we conducted targeted ablation analyses using ESM Cambrian 600M on two representative tasks, namely, *metal ion binding*, which is strongly structure-dependent, and *solubility*, which is primarily sequence-driven. Results are provided in the Supplementary Material. Comparing *h_anchor_*, *h_algn_*, and *h_enrich_* reveals clear functional specialization. The alignment representation *h_algn_*substantially improves performance on metal ion binding relative to the original sequence embedding (0.793 against 0.768 of AUCROC), while degrading performance on solubility (0.717 against 0.772 of AUCROC). This behavior confirms that the alignment subspace captures structure-related signals that are beneficial for structure-sensitive tasks but may interfere with sequence-dominated ones. In contrast, the enriched representation *h_enrich_* achieves improved performance on metal ion binding (0.771 of AUCROC) while maintaining solubility performance close to the original embedding (0.756 of AUCROC), demonstrating that residual combination enables controlled integration of structural and semantic information. The anchor representation *h_anchor_* performs slightly below the original sequence embedding, with 0.769 of AUCROC on metal ion binding and 0.751 of AUCROC on solubility, respectively, which can be attributed to dimensional compression rather than multimodal effects.

Next, we evaluated structure-only and dynamics-only variants of the model, with the results in Supplementary Material. In this ablation, we train ProtEnrich from scratch using ESM Cambrian 600M as sequence input feature, generating a model with only structural and one model with only dynamical alignment. Performance varied across tasks, with dynamical information contributing more strongly to metal ion binding, which achieved 0.782 of AUCROC, whereas structural information was more beneficial for solubility, obtaining 0.760 of AUCROC. In most tasks, the full model, with both structural and dynamical information, achieved competitive or improved performance relative to single-modality variants, indicating complementary contributions of structure and dynamics.

We further analyzed the effect of the residual scaling parameter *α*, which controls the contribution of the alignment subspace to the enriched representation. Although *α* is learnable during fine-tuning, all experiments were conducted with the pretrained model frozen and *α* = 0.039. Sensitivity analysis, in Supplementary Material, reveals a clear trade-off between structure-sensitive and sequence-dominated tasks. For metal ion binding, performance increases with larger *α* values, rising from 0.769 of AUCROC at *α* = 0 to 0.767 of AUCROC at *α* = 0.2, indicating that stronger structural injection benefits this task. In contrast, solubility peaks at intermediate scaling and declines at higher values, dropping to 0.757 of AUCROC at *α* = 0.2, close to the baseline. The default value *α* = 0.039 provides modest gains in metal ion binding while preserving solubility.

Then, we analyse the energy. Removal of energy regularization resulted in a dramatic increase in the magnitude of the alignment subspace. The mean ratio between alignment and anchor norms increased from 0.060 to 0.977, with alignment norm rising from 1.91 to 31.07. This indicates that without explicit magnitude control, the alignment representation dominates the embedding space, violating the residual design principle.

To verify separation between latent subspaces, we computed cosine similarity between *h_anchor_*and *h_algn_* on the 10,000 proteins out-of-distribution set. The mean similarity was 0.009, comparable to the similarity obtained under random permutation, which achieved 0.010, suggesting approximate orthogonality between semantic and alignment subspaces.

Finally, we examined the representation space to assess whether the model preserves structural and dynamical organization in the learned embeddings. Specifically, we evaluated whether pair-wise similarity in structure or dynamics correlates with similarity between *h_anchor_* and *h_algn_*. To ensure an unbiased evaluation, we randomly sampled 100,000 protein pairs from a set of 10,000 proteins out-of-distribution set. As shown in Figure 3, *h_algn_* preserves the geometric organization of structural space, maintaining structurally similar proteins in close proximity, with a similar but weaker trend for dynamics due to its more complex nature. In contrast, *h_anchor_* shows no meaningful correlation with structural or dynamical similarity, indicating that it primarily captures general sequence-level properties rather than higher-order biophysical information. Quantitatively, *h_algn_* achieves a Spearman correlation coefficient of 0.66 and 0.58 with structural and dynamical similarity, respectively, compared to 0.59 and 0.25 for *h_anchor_*.

**Figure 3:**
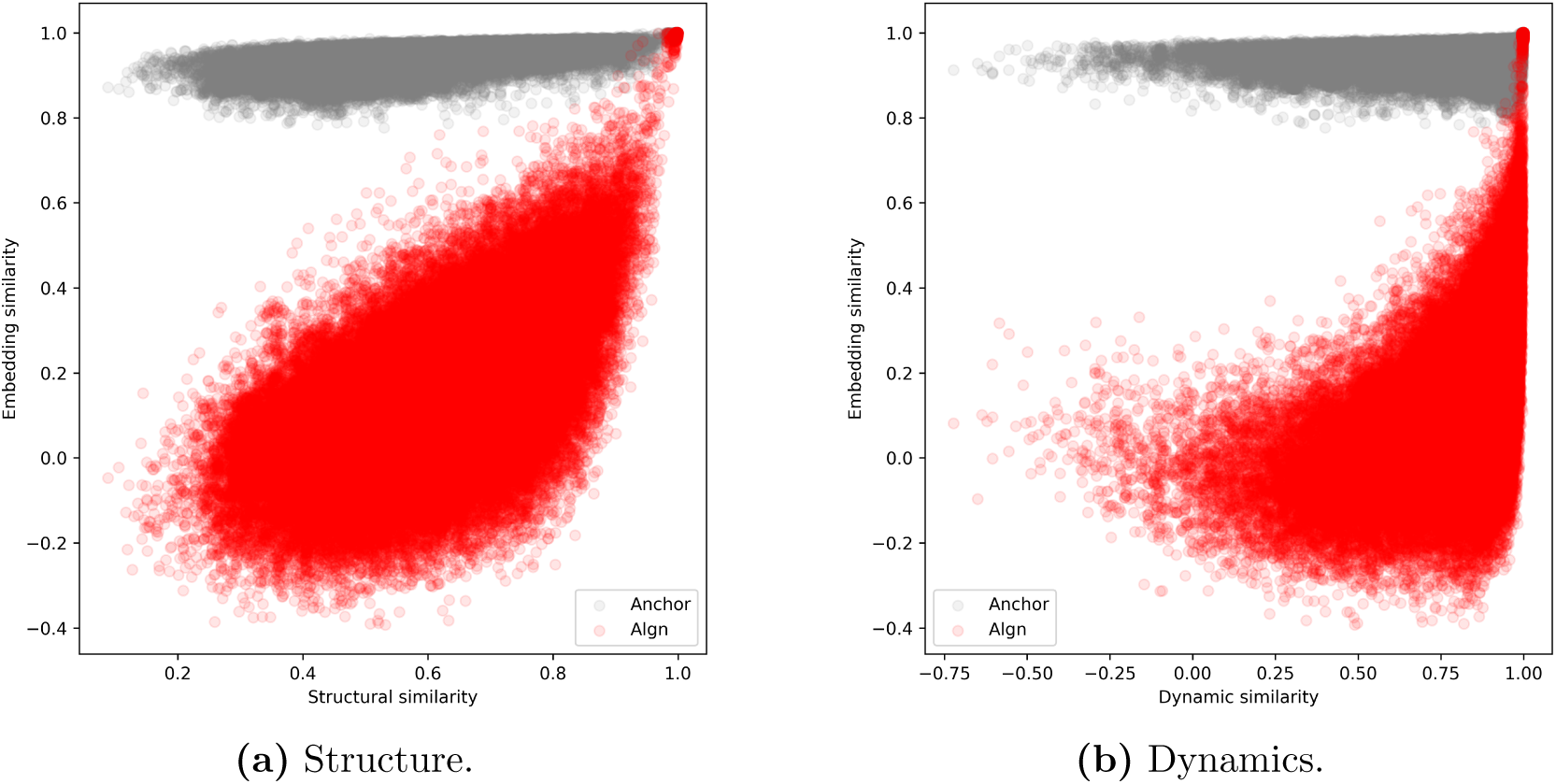
Geometric analysis of the learned latent subspaces. (a) Structure. The relationship between structural similarity and latent similarity. (b) Dynamics. The relationship between dynamical similarity and latent similarity.

## 4 Limitations and Future work

While ProtEnrich shows consistent improvements across multiple protein language models and tasks, some limitations remain. First, the framework relies on pretrained structural and dynamical representations to guide the multimodal alignment process. Although this allows the model to transfer biophysical information into sequence embeddings, the quality of the enrichment depends on the reliability of these auxiliary representations. Second, ProtEnrich operates on global protein embeddings obtained by mean pooling token representations. This design makes it easier to apply the method across different protein language models, but it may limit the ability to capture residue-level structural and dynamical relationships. Finally, this study focuses only on structural and dynamical information derived from ProstT5 and RocketSHP. Other potentially useful sources of biological information, such as domain-level annotations and protein-protein interactions, were not considered.

Future work may explore extending the residual enrichment approach to additional modalities and investigating alternative representation levels. In addition, fine-tuning strategies may further improve the ability of enriched embeddings to capture task-specific relationships.

## 5 Conclusion

In this work, we introduced *ProtEnrich*, a novel representation learning framework that injects biophysical information into protein sequence embeddings without distorting the original semantic geometry. By formalizing multimodal integration as a low-energy residual enrichment problem, we demonstrated that sequence representations can be effectively decomposed into orthogonal latent subspaces, separating sequence semantics from multimodal biophysical information. ProtEnrich implements this idea using two learned representations derived from the sequence embedding, that is, an anchor representation, that preserves sequence semantics, and an alignment representation, that captures structural and dynamical relationships.

Our extensive evaluations across eight diverse protein foundational models confirm that ProtEnrich is a model-agnostic framework capable of injecting structural and dynamical information into sequence embeddings while preserving the geometry of the original embedding space. The enriched embeddings consistently improved zero-shot remote homology detection, increasing Precision@10 and MRR by up to 0.13 and 0.11, respectively. It also provided performance gains in structure-dependent predictive tasks, such as metal ion binding (increasing up to 2.4 of AUCROC) and fluorescence prediction (up to 16% error reduction), while requiring only sequence data at inference time. Overall, ProtEnrich provides a scalable and model-agnostic strategy to bridge sequence-based representation learning with the physical constraints governing protein behavior. By enabling latent imputation of structural and conformational information directly from sequence embeddings, our framework advances large-scale computational analysis of proteins lacking experimentally resolved structures. In contrast to existing multimodal approaches that require structural inputs at inference time or modify the geometry of sequence embedding spaces, ProtEnrich enables the integration of structural and dynamical information while preserving the semantic organization of sequence representations.

## Acknowledgments

The authors thank the anonymous reviewers for their valuable suggestions. This research was supported by the NIGMS of the National Institutes of Health (NIH) under award number: R35GM153434. The content is solely the responsibility of the authors and does not necessarily represent the official views of the National Institutes of Health.

## S1. Downstream Benchmark Datasets

### Fluorescence

For this task, we used the dataset released by Frolova *et al.* [3] to predict the fluorescence intensity of mutant green fluorescent proteins [6]. The dataset consists of 21,446 training proteins, 5,362 validation proteins, and 27,217 test proteins. Fluorescence values range from 1.28 to 4.12.

### Thermostability

For thermostability prediction, we used the dataset provided by Frolova *et al.* [3], which focuses on predicting the melting temperature of human proteins [2]. The dataset contains 5,025 training proteins, 636 validation proteins, and 1,329 test proteins. Thermostability values range from 40.20 to 67.00.

### Localization

For subcellular localization, we used the dataset provided by Tan *et al.* [7] based on DeepLoc [1]. We considered two tasks: (1) binary classification to determine whether a protein is membrane-bound, and (2) multiclass classification across ten possible subcellular compartments. For the binary task, the dataset includes 5,735 training proteins, 1,009 validation proteins, and 1,728 test proteins. For the multiclass task, the dataset contains 9,324 training proteins, 1,658 validation proteins, and 2,742 test proteins.

### Metal Ion Binding

For this task, we used the dataset released by Frolova *et al.* [3] to predict whether a protein contains a metal ion binding site [4]. The dataset consists of 5,066 training proteins, 662 validation proteins, and 665 test proteins.

### Protein-Protein Interaction

For protein-protein interaction prediction, we used the dataset provided by Frolova *et al.* [3], aiming to determine whether two human proteins interact [8]. The dataset includes 5,131 proteins forming 26,317 training interactions, 215 proteins with 234 validation interactions, and 180 proteins with 180 test interactions.

### Solubility

For solubility prediction, we used the dataset introduced by Khurana *et al.* [5] to predict whether a protein is soluble. The dataset contains 62,478 training proteins, 6,942 validation proteins, and 1,999 test proteins.

### Protein Function Prediction

We constructed a protein function prediction dataset following the CAFA evaluation protocol. Data were collected from UniProt, including only proteins with manually curated Gene Ontology (GO) annotations. To ensure robust generalization, we applied homology filtering such that sequence similarity between training, validation, and test sets was below 30%. The complete dataset comprises 114,369 proteins. Function prediction is formulated as a multilabel classification task across the three Gene Ontology ontologies: Biological Process (BP), Cellular Component (CC), and Molecular Function (MF).

- **Biological Process (BP):** 80,185 training proteins, 10,127 validation proteins, and 9,910 test proteins, with 595 possible GO terms.
- **Cellular Component (CC):** 85,314 training proteins, 10,753 validation proteins, and 10,467 test proteins, with 560 possible GO terms.
- **Molecular Function (MF):** 75,072 training proteins, 9,295 validation proteins, and 9,235 test proteins, with 550 possible GO terms.

## S2. Zero-Shot Retrieval Results

Table S1 presents the detailed results for remote homology considering original embeddings against enrich embeddings *h_enrich_*obtained from ProtEnrich.

**Table S1:** Zero-shot retrieval performance on the remote homology benchmark. Metrics are reported at fold, superfamily, and family levels. A@10 is Accuracy at 10, P@10 is Precision at 10, R@10 is Recall at 10, and MRR is Mean Reciprocal Rank.

| Model | Embedding | Fold |  |  |  | Superfamily |  |  |  | Family |  |  |  |
| --- | --- | --- | --- | --- | --- | --- | --- | --- | --- | --- | --- | --- | --- |
|  |  | A@10 | P@10 | R@10 | MRR | A@10 | P@10 | R@10 | MRR | A@10 | P@10 | R@10 | MRR |
| ESM C 600M | Original | 0.822 | 0.366 | 0.096 | 0.721 | 0.806 | 0.349 | 0.132 | 0.710 | 0.801 | 0.357 | 0.212 | 0.721 |
|  | Enrich | 0.892 | 0.459 | 0.129 | 0.813 | 0.884 | 0.445 | 0.178 | 0.808 | 0.886 | 0.454 | 0.288 | 0.816 |
| ESM1b | Original | 0.972 | 0.752 | 0.228 | 0.955 | 0.972 | 0.735 | 0.312 | 0.954 | 0.973 | 0.691 | 0.479 | 0.950 |
|  | Enrich | 0.978 | 0.810 | 0.256 | 0.965 | 0.978 | 0.796 | 0.351 | 0.965 | 0.978 | 0.741 | 0.532 | 0.963 |
| ESM2 T36 | Original | 0.965 | 0.669 | 0.201 | 0.942 | 0.962 | 0.656 | 0.277 | 0.939 | 0.963 | 0.634 | 0.429 | 0.937 |
|  | Enrich | 0.978 | 0.769 | 0.241 | 0.965 | 0.976 | 0.757 | 0.332 | 0.964 | 0.977 | 0.710 | 0.502 | 0.958 |
| ProtBERT | Original | 0.837 | 0.391 | 0.093 | 0.746 | 0.820 | 0.374 | 0.129 | 0.733 | 0.808 | 0.369 | 0.204 | 0.733 |
|  | Enrich | 0.913 | 0.517 | 0.134 | 0.854 | 0.905 | 0.501 | 0.184 | 0.848 | 0.901 | 0.490 | 0.294 | 0.846 |
| Ankh 3 XL | Original | 0.978 | 0.678 | 0.215 | 0.960 | 0.978 | 0.668 | 0.297 | 0.959 | 0.977 | 0.658 | 0.467 | 0.957 |
|  | Enrich | 0.985 | 0.727 | 0.235 | 0.973 | 0.985 | 0.720 | 0.324 | 0.973 | 0.985 | 0.702 | 0.506 | 0.971 |
| ProtT5 | Original | 0.980 | 0.729 | 0.227 | 0.962 | 0.979 | 0.715 | 0.312 | 0.960 | 0.980 | 0.683 | 0.481 | 0.954 |
|  | Enrich | 0.987 | 0.770 | 0.243 | 0.975 | 0.986 | 0.760 | 0.336 | 0.974 | 0.987 | 0.721 | 0.519 | 0.970 |
| ProGen2 | Original | 0.832 | 0.443 | 0.110 | 0.753 | 0.818 | 0.428 | 0.147 | 0.747 | 0.815 | 0.417 | 0.237 | 0.749 |
|  | Enrich | 0.884 | 0.561 | 0.150 | 0.825 | 0.873 | 0.545 | 0.200 | 0.821 | 0.874 | 0.515 | 0.318 | 0.820 |
| CARP 640M | Original | 0.967 | 0.724 | 0.204 | 0.941 | 0.963 | 0.692 | 0.275 | 0.936 | 0.957 | 0.620 | 0.411 | 0.919 |
|  | Enrich | 0.970 | 0.749 | 0.217 | 0.943 | 0.968 | 0.720 | 0.294 | 0.938 | 0.965 | 0.646 | 0.438 | 0.922 |

Table S2 presents the detailed results for zero-shot based on similarity of structure and dynamics considering original embeddings against enrich embeddings *h_enrich_* obtained from ProtEnrich.

**Table S2:** Zero-shot retrieval performance for structural and dynamical similarity. A@10 is Accuracy at 10, P@10 is Precision at 10, R@10 is Recall at 10, and MRR is Mean Reciprocal Rank.

| Model | Embedding | Dynamics |  |  |  | Structure |  |  |  |
| --- | --- | --- | --- | --- | --- | --- | --- | --- | --- |
|  |  | A@10 | P@10 | R@10 | MRR | A@10 | P@10 | R@10 | MRR |
| ESM C 600M | Original | 0.856 | 0.356 | 0.036 | 0.714 | 0.943 | 0.474 | 0.047 | 0.802 |
|  | Enrich | 0.868 | 0.385 | 0.039 | 0.732 | 0.951 | 0.536 | 0.054 | 0.831 |
| ESM1b | Original | 0.894 | 0.412 | 0.041 | 0.756 | 0.970 | 0.595 | 0.060 | 0.858 |
|  | Enrich | 0.902 | 0.431 | 0.043 | 0.765 | 0.975 | 0.624 | 0.062 | 0.866 |
| ESM2 T36 | Original | 0.872 | 0.395 | 0.040 | 0.739 | 0.949 | 0.543 | 0.054 | 0.838 |
|  | Enrich | 0.885 | 0.415 | 0.042 | 0.752 | 0.961 | 0.583 | 0.058 | 0.853 |
| ProtBERT | Original | 0.844 | 0.323 | 0.032 | 0.691 | 0.939 | 0.451 | 0.045 | 0.791 |
|  | Enrich | 0.875 | 0.374 | 0.037 | 0.724 | 0.959 | 0.537 | 0.054 | 0.831 |
| Ankh 3 XL | Original | 0.899 | 0.418 | 0.042 | 0.758 | 0.982 | 0.652 | 0.065 | 0.874 |
|  | Enrich | 0.893 | 0.419 | 0.042 | 0.755 | 0.976 | 0.638 | 0.064 | 0.869 |
| ProtT5 | Original | 0.899 | 0.424 | 0.042 | 0.755 | 0.983 | 0.664 | 0.066 | 0.878 |
|  | Enrich | 0.894 | 0.426 | 0.043 | 0.757 | 0.976 | 0.653 | 0.065 | 0.874 |
| ProGen2 | Original | 0.850 | 0.361 | 0.036 | 0.706 | 0.933 | 0.491 | 0.049 | 0.797 |
|  | Enrich | 0.883 | 0.387 | 0.039 | 0.730 | 0.954 | 0.548 | 0.055 | 0.828 |
| CARP 640M | Original | 0.910 | 0.415 | 0.042 | 0.758 | 0.977 | 0.609 | 0.061 | 0.859 |
|  | Enrich | 0.911 | 0.420 | 0.042 | 0.760 | 0.979 | 0.622 | 0.062 | 0.862 |

## S3. Latent Feature Imputation Results

We compared seven input configurations for latent feature imputation on filtered metal ion binding dataset:

1. Sequence only: original sequence embedding.
2. Sequence + Structure (original): concatenation of the original sequence embedding with the original structural embedding.
3. Sequence + Structure (generated): concatenation of the original sequence embedding with the structure embedding generated by ProtEnrich.
4. Sequence + Dynamics (original): concatenation of the original sequence embedding with the original dynamical embedding.
5. Sequence + Dynamics (generated): concatenation of the original sequence embedding with the dynamics embedding generated by ProtEnrich.
6. Sequence + Structure (original) + Dynamics (original): concatenation of sequence with both original structure and dynamics embeddings.
7. Sequence + Structure (generated) + Dynamics (generated): concatenation of sequence with both generated structure and generated dynamics.

The results are presented in Table S3.

**Table S3:** Metal ion binding classification. We compare original sequence embeddings alone against concatenations with original or generated structural and dynamical embeddings.

| Configuration | ESM C | ProtBERT | ProtT5 | Ankh | ESM2 T36 | ESM1b | ProGen2 | CARP 640M |
| --- | --- | --- | --- | --- | --- | --- | --- | --- |
| Seq only | 0.751 | 0.732 | 0.802 | 0.779 | 0.774 | 0.757 | 0.767 | 0.770 |
| Seq + Str (orig) | 0.821 | 0.793 | 0.831 | 0.814 | 0.803 | 0.765 | 0.770 | 0.765 |
| Seq + Str (gen) | 0.786 | 0.753 | 0.817 | 0.768 | 0.777 | 0.749 | 0.788 | 0.766 |
| Seq + Dyn (orig) | 0.777 | 0.736 | 0.806 | 0.752 | 0.796 | 0.777 | 0.771 | 0.777 |
| Seq + Dyn (gen) | 0.742 | 0.713 | 0.799 | 0.764 | 0.788 | 0.767 | 0.768 | 0.773 |
| Seq + Str (orig) + Dyn (orig) | 0.816 | 0.792 | 0.829 | 0.817 | 0.809 | 0.772 | 0.783 | 0.763 |
| Seq + Str (gen) + Dyn (gen) | 0.789 | 0.740 | 0.815 | 0.789 | 0.788 | 0.759 | 0.765 | 0.768 |

## S4. Comparison with Structure-Aware Pretraining

Table S4 reports zero-shot remote homology retrieval at fold, superfamily, and family levels between the original ESM2 T36 embeddings, ProtEnrich, and ISM.

**Table S4:** Zero-shot retrieval performance between the original ESM2 T36 embeddings, ProtEnrich, and ISM on the remote homology benchmark. Metrics are reported at fold, superfamily, and family levels. A@10 is Accuracy at 10, P@10 is Precision at 10, R@10 is Recall at 10, and MRR is Mean Reciprocal Rank.

| Model | Embedding | Fold |  |  |  | Superfamily |  |  |  | Family |  |  |  |
| --- | --- | --- | --- | --- | --- | --- | --- | --- | --- | --- | --- | --- | --- |
|  |  | A@10 | P@10 | R@10 | MRR | A@10 | P@10 | R@10 | MRR | A@10 | P@10 | R@10 | MRR |
| ESM2 T36 | Original | 0.965 | 0.669 | 0.201 | 0.942 | 0.962 | 0.656 | 0.277 | 0.939 | 0.963 | 0.634 | 0.429 | 0.937 |
|  | ProtEnrich | 0.978 | 0.769 | 0.241 | 0.965 | 0.976 | 0.757 | 0.332 | 0.964 | 0.977 | 0.710 | 0.502 | 0.958 |
|  | ISM | 0.972 | 0.693 | 0.209 | 0.951 | 0.969 | 0.671 | 0.283 | 0.947 | 0.966 | 0.639 | 0.436 | 0.944 |

Table S5 presents zero-shot structural similarity retrieval results between the original ESM2 T36 embeddings, ProtEnrich, and ISM.

**Table S5:** Zero-shot retrieval performance for structural similarity between the original ESM2 T36 embeddings, ProtEnrich, and ISM. A@10 is Accuracy at 10, P@10 is Precision at 10, R@10 is Recall at 10, and MRR is Mean Reciprocal Rank.

| Model | Embedding | Structure |  |  |  |
| --- | --- | --- | --- | --- | --- |
|  |  | A@10 | P@10 | R@10 | MRR |
| ESM2 T36 | Original | 0.949 | 0.543 | 0.054 | 0.838 |
|  | ProtEnrich | 0.961 | 0.583 | 0.058 | 0.853 |
|  | ISM | 0.988 | 0.670 | 0.067 | 0.884 |

The downstream performance comparison between the original ESM2 T36 embeddings, ProtEnrich, and ISM on fluorescence and metal ion binding is summarized in Table S6.

**Table S6:** Downstream comparison between the original ESM2 T36 embeddings, ProtEnrich, and ISM on fluorescence (RMSE, lower is better) and metal ion binding (AUC, higher is better). Results are reported as mean *±* standard deviation across runs.

| Model | Embedding | Fluorescence ↓ | MIB ↑ |
| --- | --- | --- | --- |
| ESM2 T36 | Original | 0.871 ± 0.027 | 0.809 ± 0.017 |
|  | ProtEnrich | 0.924 ± 0.021 | 0.833 ± 0.009 |
|  | ISM | 0.868 ± 0.051 | 0.821 ± 0.008 |

## S5. Ablation Study Results

Table S7 presents the results for the original embeddings, *h_enrich_*, *h_algn_*, and *h_anchor_* based on ESM Cambrian 600M for metal ion binding (MIB) and solubility (Sol).

**Table S7:** Ablation study comparing the original embeddings, *h_enrich_*, *h_algn_*, and *h_anchor_* based on ESM Cambrian 600M on metal ion binding (MIB) and solubility (Sol).

| Embedding | MIB | Sol |
| --- | --- | --- |
| Original | $0.768 \pm 0.014$ | $0.772 \pm 0.005$ |
| $h_{enrich}$ | $0.771 \pm 0.013$ | $0.756 \pm 0.008$ |
| $h_{anchor}$ | $0.769 \pm 0.013$ | $0.751 \pm 0.005$ |
| $h_{align}$ | $0.793 \pm 0.009$ | $0.717 \pm 0.005$ |

Table S8 presents the results for the *α* value using ESM Cambrian 600M embedings on metal ion binding (MIB) and solubility (Sol).

**Table S8:** Ablation study comparing the *α* value using ESM Cambrian 600M embedings on metal ion binding (MIB) and solubility (Sol).

| Alpha | MIB | Sol |
| --- | --- | --- |
| 0.000 | $0.769 \pm 0.015$ | $0.751 \pm 0.005$ |
| 0.039 | $0.771 \pm 0.013$ | $0.756 \pm 0.008$ |
| 0.050 | $0.768 \pm 0.010$ | $0.754 \pm 0.007$ |
| 0.100 | $0.764 \pm 0.013$ | $0.746 \pm 0.006$ |
| 0.150 | $0.765 \pm 0.010$ | $0.760 \pm 0.004$ |
| 0.200 | $0.767 \pm 0.012$ | $0.757 \pm 0.006$ |

Table S9 presents the results for full model (using structural and dynamical representation), structure-only, and dynamics-only for downstream tasks.

**Table S9:**
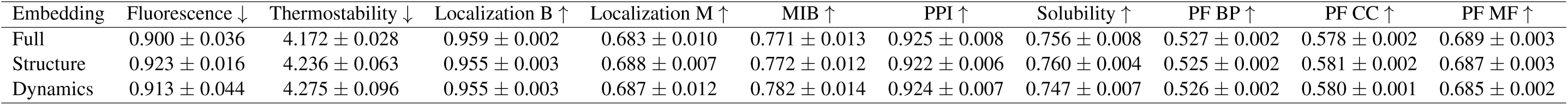
Downstream performance (mean *±* standard deviation) for the enriched using structure and dynamics (full), structure-only, and dynamics-ony embeddings. Localization B and Localization M denote binary and multiclass localization, respectively. MIB refers to metal ion binding, and PPI to protein-protein interaction. PF stands for protein function prediction and is divided into three Gene Ontology categories: BP (Biological Process), CC (Cellular Component), and MF (Molecular Function). For regression tasks, lower values indicate better performance, whereas for classification tasks, higher values indicate better performance.

| Embedding | Fluorescence ↓ | Thermostability ↓ | Localization B ↑ | Localization M ↑ | MIB ↑ | PPI ↑ | Solubility ↑ | PF BP ↑ | PF CC ↑ | PF MF ↑ |
| --- | --- | --- | --- | --- | --- | --- | --- | --- | --- | --- |
| Full | 0.900 ± 0.036 | 4.172 ± 0.028 | 0.959 ± 0.002 | 0.683 ± 0.010 | 0.771 ± 0.013 | 0.925 ± 0.008 | 0.756 ± 0.008 | 0.527 ± 0.002 | 0.578 ± 0.002 | 0.689 ± 0.003 |
| Structure | 0.923 ± 0.016 | 4.236 ± 0.063 | 0.955 ± 0.003 | 0.688 ± 0.007 | 0.772 ± 0.012 | 0.922 ± 0.006 | 0.760 ± 0.004 | 0.525 ± 0.002 | 0.581 ± 0.002 | 0.687 ± 0.003 |
| Dynamics | 0.913 ± 0.044 | 4.275 ± 0.096 | 0.955 ± 0.003 | 0.687 ± 0.012 | 0.782 ± 0.014 | 0.924 ± 0.007 | 0.747 ± 0.007 | 0.526 ± 0.002 | 0.580 ± 0.001 | 0.685 ± 0.002 |

